# Saving the Devils is in the details: Tasmanian Devil facial tumor disease can be eliminated with interventions

**DOI:** 10.1101/2022.04.03.486872

**Authors:** Brian Drawert, Sean Matthew, Megan Powell, Bryan Rumsey

## Abstract

Tasmanian Devils facial tumor disease is severely impacting the population of this wild animal. We developed a computational model of the population of Tasmanian Devils, and the change induced by DFTD. We use this model to test possible intervention strategies Tasmanian conservationists could do. We investigate bait drop vaccination programs, diseased animal removals programs, and evolution of natural immunity. We conclude that a combination of intervention strategies gives the most favorable outcome.

An additional goal of this paper is for reproducibility of our results. Our StochSS software platform features the ability to share and reproduce the computational notebooks that created all of the results in the paper. We endeavor that all readers should be able to reproduce our results with minimum effort.

## 1 Introduction

Devil facial tumor disease (DFTD) has been ravaging the Tasmanian devil population since it was first seen in the wild in 1996. There is an estimated 80% decline in pre-DFTD population in parts of Tasmania (McCallum et al., 2007). As the apex predator of the Tasmanian ecosystem, it’s precipitous decrease in population numbers has lead to notable changes in the species ecosystem including a significant increase in feral cat populations and decrease in quoll populations (Fancourt et al., 2015; Hollings et al., 2014, 2016). Additionally, the disease, spread through biting primarily during mating, is disproportionately affecting devils with higher fitness, causing a decrease in the overall fitness of the devil population (Hohenlohe, 2017). Some natural resistance to the disease as well as tumor regression has been seen, but it is unclear if this resistance alone is enough to help the wild population rebound towards pre-DFTD numbers (Hohenlohe et al., 2019; Pye et al., 2016; Wright et al., 2017; Margres et al., 2020; Hamede et al., 2021). Without human intervention, it is possible the devil will go extinct in the wild (McCallum et al., 2007) and current captive and wild insurance populations will be necessary to keep the species alive (Rout et al., 2017). Selective culling of diseased animals has been considered in the past but models show it would not be effective alone (Beeton and McCallum, 2011) but DFTD vaccine development has been promising in recent years (Flies et al., 2020; Tovar et al., 2017; Owen and Siddle, 2019).

In this paper we consider the effectiveness of bait drop or trap-vaccinate-release wildlife vaccines either as a single intervention or combined with selective culling of only highly diseased animals who are no longer reproducing. We find that successful elimination of the disease is the only way to have long term recovery of the devil population to pre-DFTD numbers. Additionally, we explore the preliminary data evidence that devils may be developing some natural immunity to the disease and how much immunity would need to develop to allow for a sustainable population without human intervention (Hamede et al., 2021).

## 2 Background

We develop a modified SEIV model and calibrate to the population data presented in (Cunningham et al., 2021). Through the parameter estimation process we find there is an additional class of highly diseased devils who are no longer successfully reproducing who we document as in the Diseased class (Hamilton et al., 2019, 2020). Additionally, we are able to estimate the actual start of DFTD to be 1993. After successfully parameterizing the model with and without the disease, we consider the population trajectory if

- A successful vaccine and bait drop process is developed and implemented.
- Highly diseased animals in the Diseased class are culled to remove them from the population.
- Devils develop natural resistance to DFTD that slows down movement between Susceptible, Exposed, Infected, and Diseased classes.
- None of the above occur.

We find successful strategies include vaccination alone, development of high levels of natural immunity, vaccination plus culling of highly diseased animals. Our findings are consistent with previous modeling efforts that conclude selective culling alone will not be effective (Beeton and McCallum, 2011) and support models that show vaccine efforts may be the most effective strategy (Bruno et al., 2017; Bobbitt et al., 2020). We find that devils would need to develop at least a 60% natural immunity to DFTD for the disease to have a high probability of elimination. Therefore, our model supports that a vaccine strategy, even if it is not sufficient to eliminate the disease, may help the devils maintain a sustainable population while such natural immunity may be developing.

## 3 Model Description

We developed a stochastic model whose diagram is shown in Figure 2. Equations 1-6 describe the base population model without any disease. Adult devils produce juvenile devils at a rate *K*_*birth*_ and juvenile devils become adult devils at a rate *K*_*mature*_. Juvenile devils mature into adult devils. Both juveniles and adults have non-disease death rates of *D*_*devil*_ and *D*_*juvenile*_ respectively as well as a combined death rate due to overcrowding of *D*_*overcrowding*_. The rate parameters in Equations. 1-6 were fit to data from (Cunningham et al., 2021) assuming a carrying capacity of 61322 devils for the whole ecosystem.

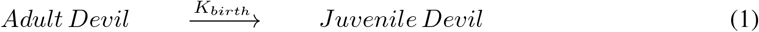

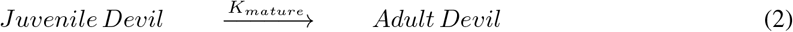

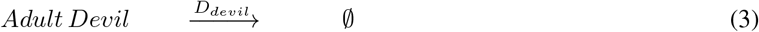

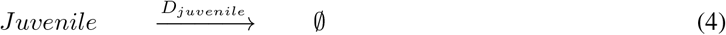

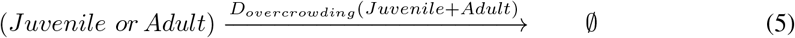

**Figure 1:**
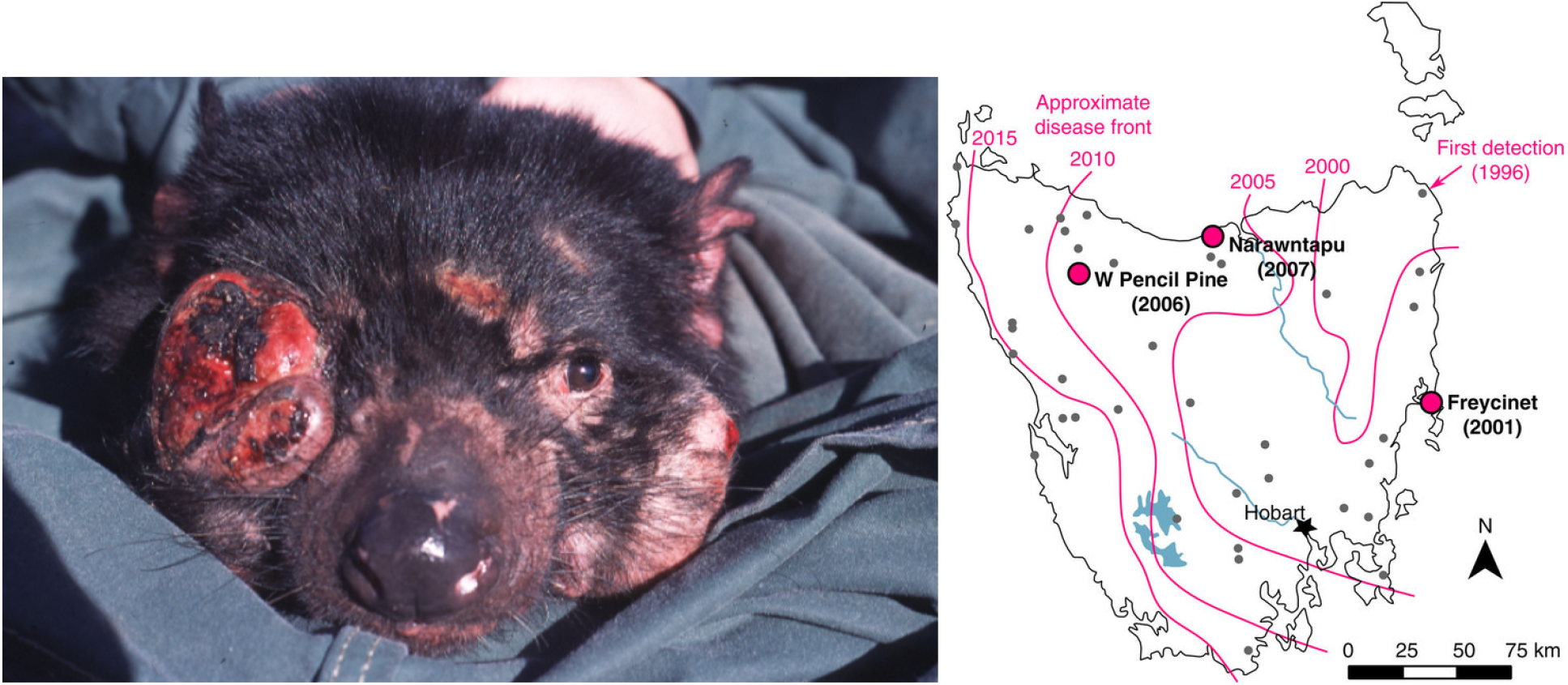
(a) Devil facial tumour disease causes tumours to form in and around the mouth, (Photo: Menna Jones) adapted from: (McCallum and Jones, 2006). (b) Spread of DFTD across Tasmania, adapted from:(Epstein et al., 2016).

**Figure 2:**
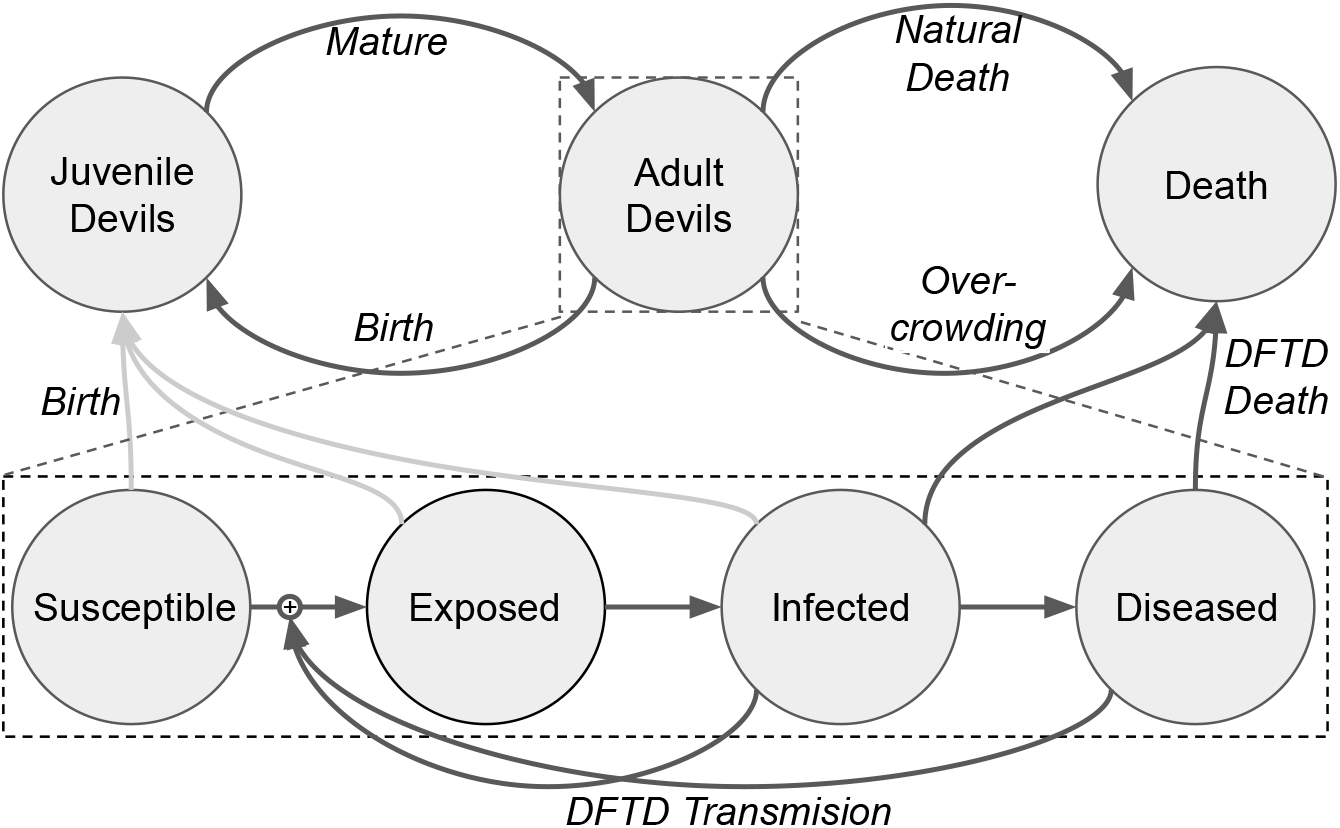
Diagram of our model. We sub-divide “Adult Devil” into four stages of the disease: Susceptible (devils without DFTD), Exposed (Devils who have been exposed to, but are not spreading DFTD), Infected (Devil who are spreading DFTD), and Diseased (Devils who are spreading DFTD and are heavily impacted by it). Note, we assume that Diseased devils do not reproduce.

To model the disease we sub-divide “Adult Devil” into four stages of the disease: **Susceptible** (devils without DFTD), **Exposed** (Devils who have been exposed to, but are not spreading DFTD), **Infected** (Devil who are spreading DFTD), and **Diseased** (Devils who are spreading DFTD and are heavily impacted by it), as shown in equations 6-11. Note, we assume that Diseased devils do not successfully reproduce due to their advanced diseased state. When Susceptible devils interact with either Infected or Diseased devils, the outcome is Exposed and Infected or Diseased devils respectively at rates *I*_*infected*_ and *I*_*diseased*_. Exposed devils move to the Infected class after the incubation period at a rate 1*/T*_*incubation*_ and similarly Infected devils move to the Diseased after the disease progresses at a rate 1*/T*_*progression*_. Both Infected and Diseased Devils die at a rate *D*_*infected*_ and *D*_*diseased*_.

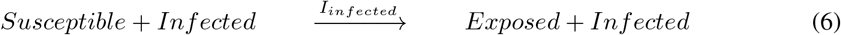

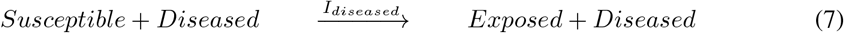

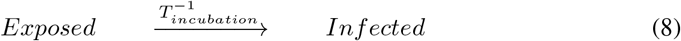

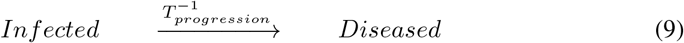

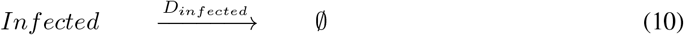

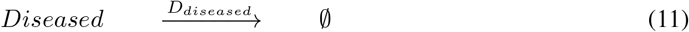

The model describe by Equations 1-11 were fit to data presented in (Cunningham et al., 2021) and were chosen based on error minimization, visual fit of the model, and consistency with field data on the disease. Figure 3A shows the reported total population data as well as our fitted model, extrapolating the devil population if DFTD had not existed. Figure 3B shows the reported total population and our fitted total population with DFTD as well as the sub-classes of devils in the model. Table 1 shows average value for the fitted parameter values for the disease. The average time from first exposure to the disease to death from the disease (*T*_*incubation*_ + *T*_*progression*_ + 1*/D*_*diseased*_) is approximately 24 months. Devils may die from other causes before this two years. While total time from exposure to death may be shorter in individual devils, we are choosing these parameters that are both biologically feasible and fit the Cunningham (Cunningham et al., 2021) data as best as possible.

**Table 1:** Model Parameters

| Parameter | Value | units |
| --- | --- | --- |
| Initial juvenile population | 16165 |  |
| Initial adult population | 18450 |  |
| $K_{birth}$ | 0.055 | Devils<sup>-1</sup> months<sup>-1</sup> |
| $K_{mature}$ | 0.04 | Devils<sup>-1</sup> months<sup>-1</sup> |
| $D_{juvenile}$ | 0.007 | Devils<sup>-1</sup> months<sup>-1</sup> |
| $D_{susceptible}$ | 0.02335 | Devils<sup>-1</sup> months<sup>-1</sup> |
| $D_{overcrowding}$ | 2.3e-07 | Devils<sup>-2</sup> months<sup>-1</sup> |
| $I_{infected}$ | 1.0e-05 | Devils<sup>-1</sup> months<sup>-1</sup> |
| $I_{diseased}$ | 3.84e-05 | Devils<sup>-1</sup> months<sup>-1</sup> |
| $T_{incubation}$ | 10.25 | months Devils |
| $T_{progression}$ | 10.74 | months Devils |
| $D_{infected}$ | 0.022609 | Devils<sup>-1</sup> months<sup>-1</sup> |
| $D_{diseased}$ | 0.29017 | Devils<sup>-1</sup> months<sup>-1</sup> |
| DFTD start | 100 | months (May 1993) |

**Figure 3:**
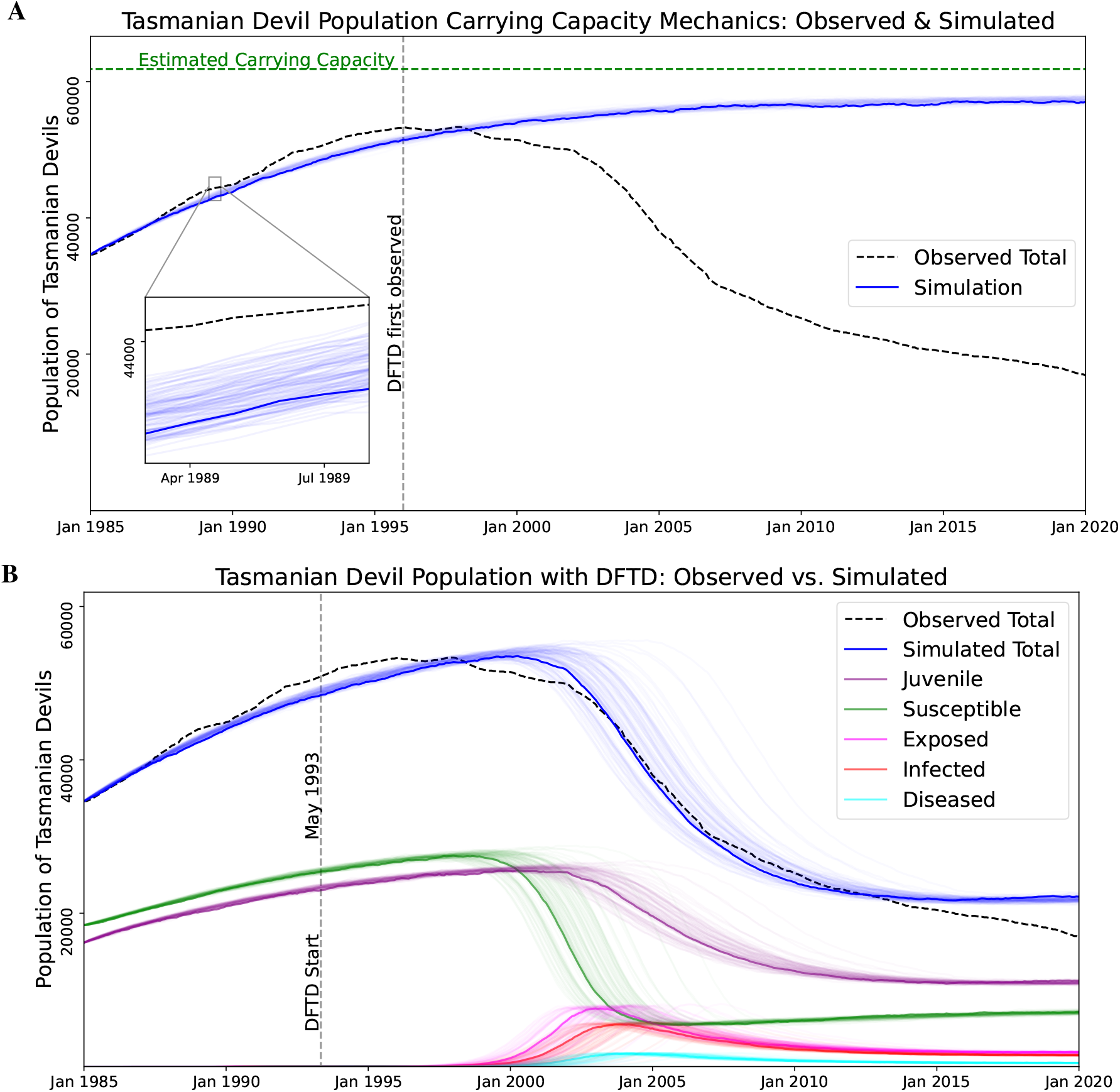
Simulations of 100 stochastic trajectories of the Tasmanian Devil population model. The first trajectory is dark, while the rest are lighter. The dashed black line represents observed Tasmanian Devil populations (Cunningham et al., 2021). **A:** Simulation of the model had DFTD not existed: simulated total population (blue lines) and observed (dashed black line). **B:** Simulation of model with DFTD: simulated total population (blue lines), simulated sub-groups (colored lines), and observed dashed black line).

## 4 Conservation Intervention Strategies

We consider how a vaccine campaign, a culling campaign or a combination of both could impact long term devil populations. We aim to find a parameter space that eliminates DFTD and allows devil populations recover to their pre-DFTD populations. Additionally, we explore how much natural immunity in what time frame would be needed for devils to sustain their population without human intervention. We show how a vaccine campaign may allow maintenance of a sustainable population while devils develop natural immunity even if the vaccine campaign is not sufficient to eliminate the disease. First we model devil populations if no interventions are present as comparison.

### 4.1 No Intervention

If no intervention is done, we see in Figure 4 that the model predicts that DFTD will become endemic and devils will maintain a population of roughly one third of their peak population. We do note Figure 4 shows juvenile devils as the largest class of devils in the endemic phase. This is troubling because juvenile devils are the most susceptible to death due to external influences such as from vehicle strikes and domestic dog attacks (Hobday et al., 2008; Jones, 2000). Our model does not account for these potentially higher death rates, therefore it is possible that the long term population would decrease more than the figure shows due to the increased proportion of the population being juveniles.

**Figure 4:**
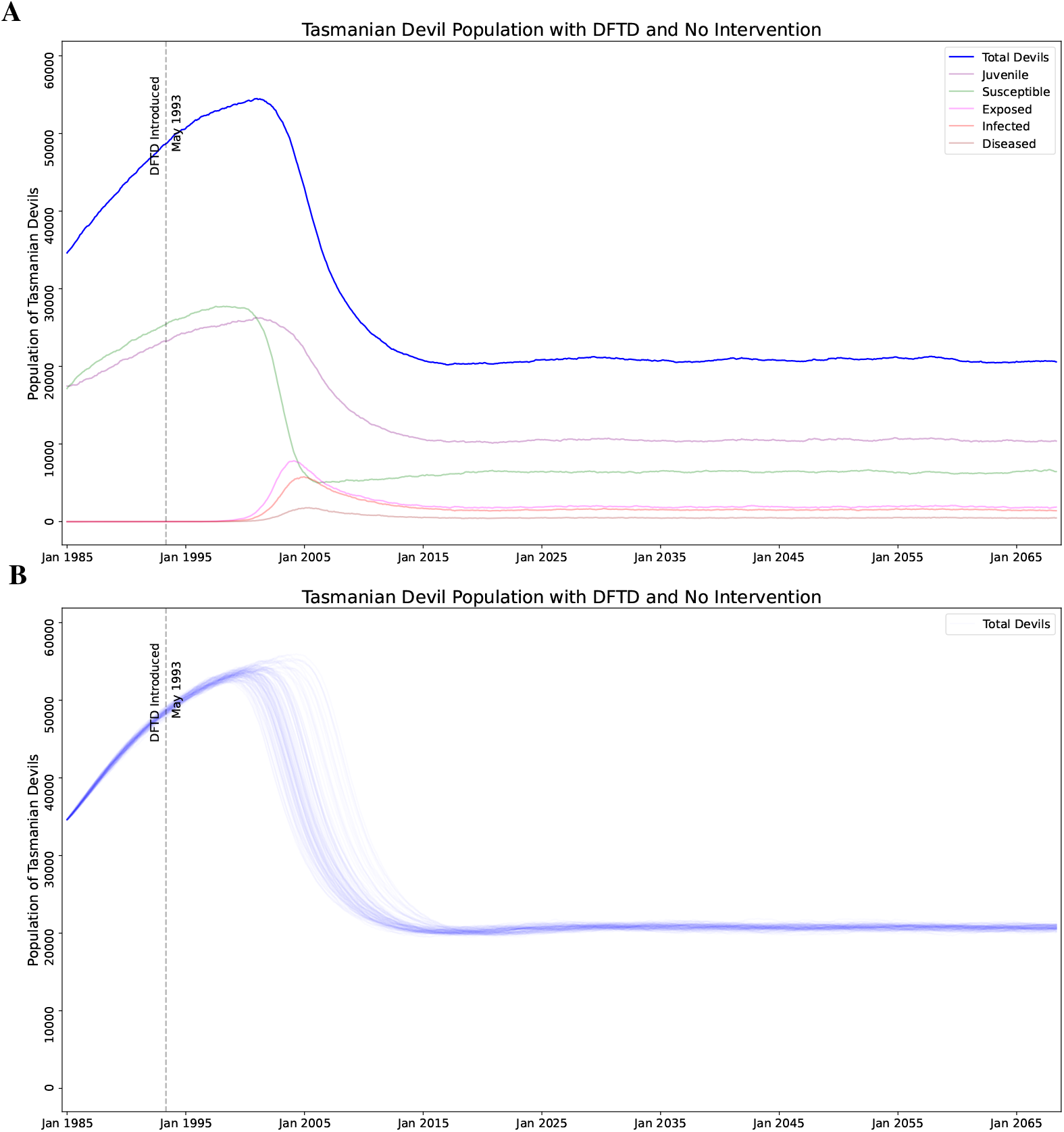
Simulation of the model with no interventions, for comparison. Note that with no human intervention DFTD become endemic and the population of devils reaches a steady-state. Also note, however, that the population of juvenile devils is larger than adult devils in this state. **A:** A single stochastic trajectory, showing all the sub-classes. **B:** 100 stochastic trajectories, showing only total devil population.

### 4.2 Evolved Natural Immunity

It has been hypothesized that the devils are becoming naturally immune through strong evolutionary selection pressure (Hamede et al., 2021) with evidence of tumor regression in rare cases (Pye et al., 2016; Wright et al., 2017; Margres et al., 2020). We model this by reducing the transmission rate of the disease by multiplying by 1.0 − *immunity*. For example, if we assume immunity is 60%, this means an individual devil will only contract the disease at 40% the rate it would had the devil no natural immunity. Due to the rarity of tumor regression, we do not model a way for devils to move backwards in the disease classes from Diseased to Infected, Infected to Exposed, or Exposed to Susceptible. This may be modeled in future efforts if more evidence of natural regression or vaccine induced regression is well documented. Figure 5 shows simulations of this program assuming that natural immunity began to develop in January 2022 growing at an exponential rate of 0.0075 to a maximum of 70% immunity. The exponential rates 0.005, 0.0075, 0.01 corresponds to approximately 23, 15, and 11 years for the devil population to gain half of the max level, respectively. We note that if devils neutrally develop a high level of immunity to the disease, they may be able to recover on their own without human intervention. We consider how high the immunity must reach for potential disease elimination by considering the maximum immunity level reached as well as when natural immunity started to develop, see equations 12-14 for more details. Figure 6 shows that at only 50% maximum immunity, elimination of DFTD is not possible, but at 70% maximum immunity, the devils may be able to eliminate the disease. There is not sufficient evidence that this level of natural immunity is achievable at all or in what time frame it would take to achieve such a high level of natural immunity.

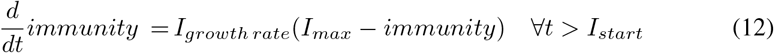

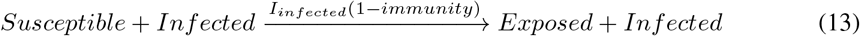

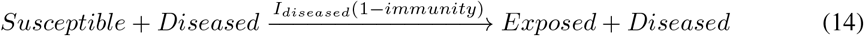

**Figure 5:**
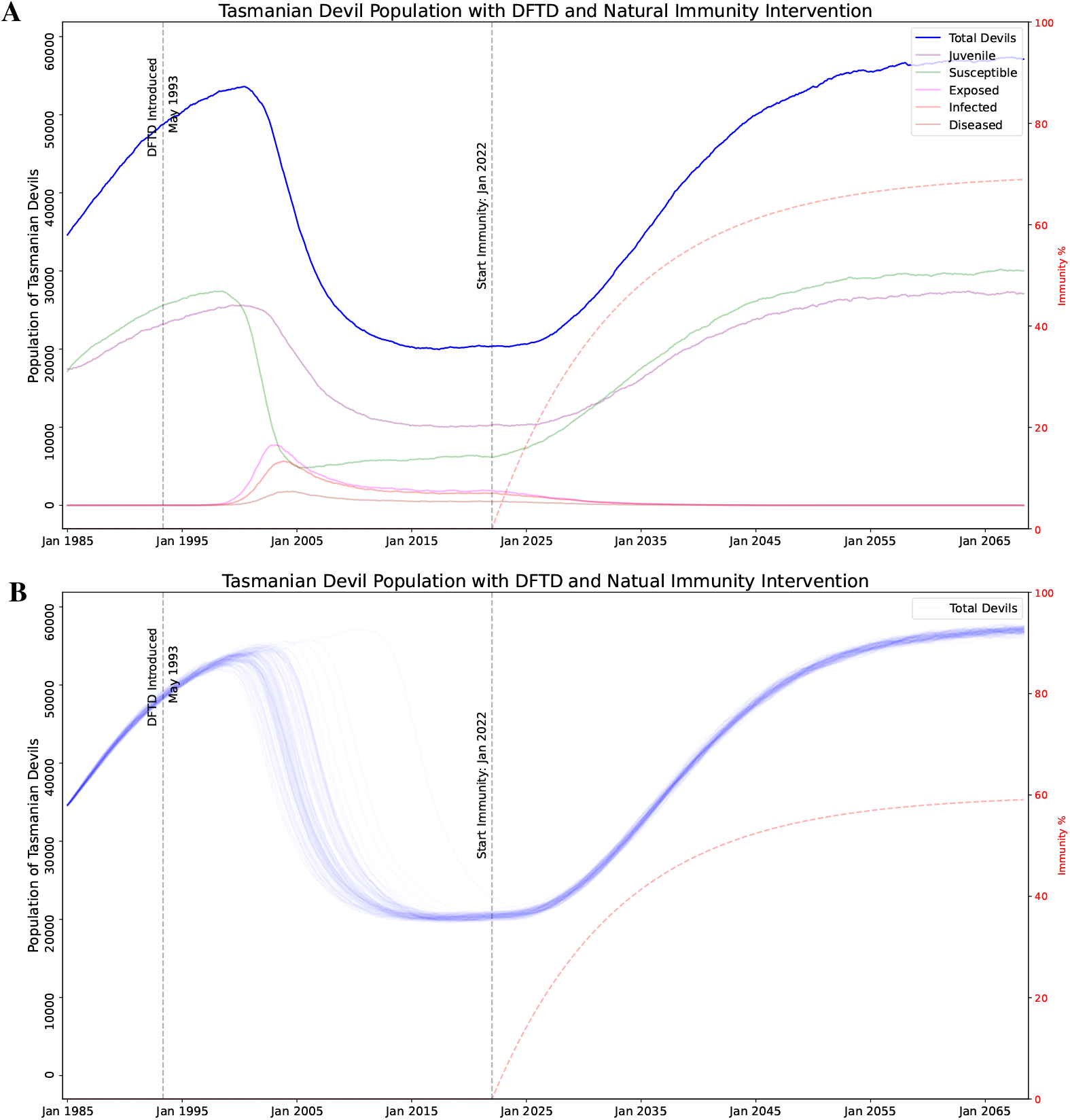
Simulation of DFTD model with naturally evolved immunity. Parameters of the program: starting Jan 2022 at 0% and growing exponentially at a rate of 0.0075 to a maximum of 70%. Even in cases where evolved natural immunity does not eliminate it, DFTD becomes endemic in the devil population and the population trends towards the carrying capacity of the environment with a more sustainable adult devil population (in contrast with the “no intervation” results). **A:** A single stochastic trajectory, showing all the sub-classes. **B:** 100 stochastic trajectories, showing only total devil population.

**Figure 6:**
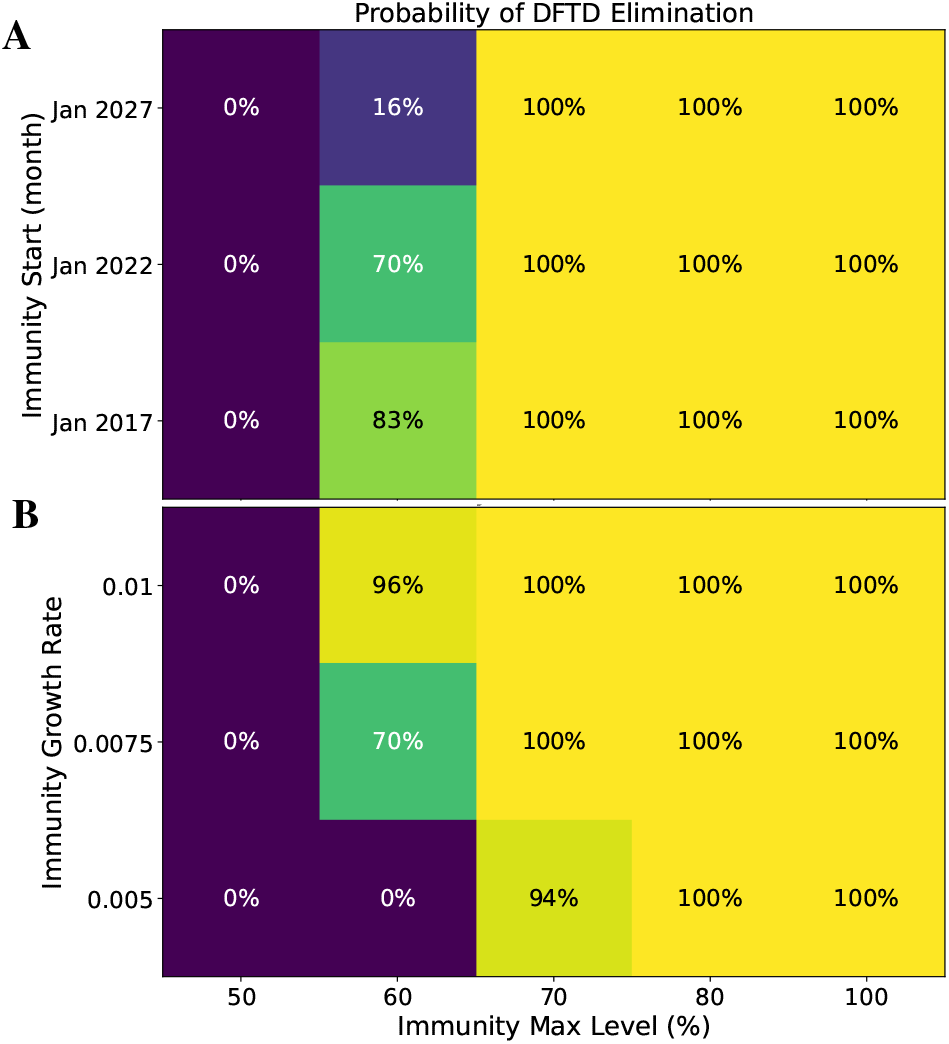
Although there is currently no evidence that Tasmanian Devils are or will develop immunity to DFTD, we explored the outcomes if such immunity evolved. Probability of eliminating DFTD with naturally evolved immunity reaching of a maximum level of 50-100% and starting to evolve between 2017 and 2027. Probability of eliminating DFTD with natural immunity, showing immunity max level versus **A:** Immunity Start, with immunity growth rate of 0.0075 (15 years to reach the half-maximum immunity level). **B:** Immunity Growth Rate, with immunity start as Jan 2022.

### 4.3 Culling

One possible intervention is culling of devils who are heavily impacted by DFTD (the “Diseased” sub-class in our model, see equation 15 and table 3). These are animals that are generally in the last 3 months of life and often die due to the disease. For our analysis, culling indicates the individuals are removed from the general population. The implementation of this intervention in the field could take many forms including a capture and quarantine type program. Figure 7 shows simulations of this program. We note that while there will be small rise in devil populations with a culling program, the disease will quickly return to its current endemic state after the conclusion of a culling program. Additionally, Figure 8 shows a 0% probability of eliminating DFTD using culling strategies alone with a culling program from 3-11 years and culling anywhere from 25% to 75% of the Diseased animals. Therefore, our model supports that culling alone is not a viable long term strategy.

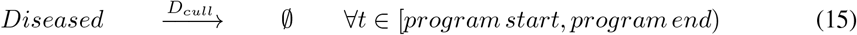

**Table 2:** Parameters for natural immunity

| Parameter | Range | Description |
| --- | --- | --- |
| Start ($I_{start}$) | 2017 → 2027 | Date when Immunity begins |
| Growth Rate ($I_{growth\ rate}$) | 0.005 → 0.01 | Exponential growth rate of immunity |
| Max Level ($I_{max}$) | 0% → 100% | Maximum level of immunity |

**Table 3:**
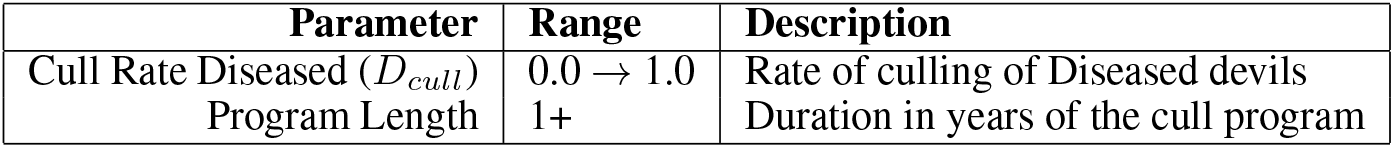
Parameters for culling intervention

| Parameter | Range | Description |
| --- | --- | --- |
| Cull Rate Diseased ($D_{cull}$) | 0.0 → 1.0 | Rate of culling of Diseased devils |
| Program Length | 1+ | Duration in years of the cull program |

**Figure 7:**
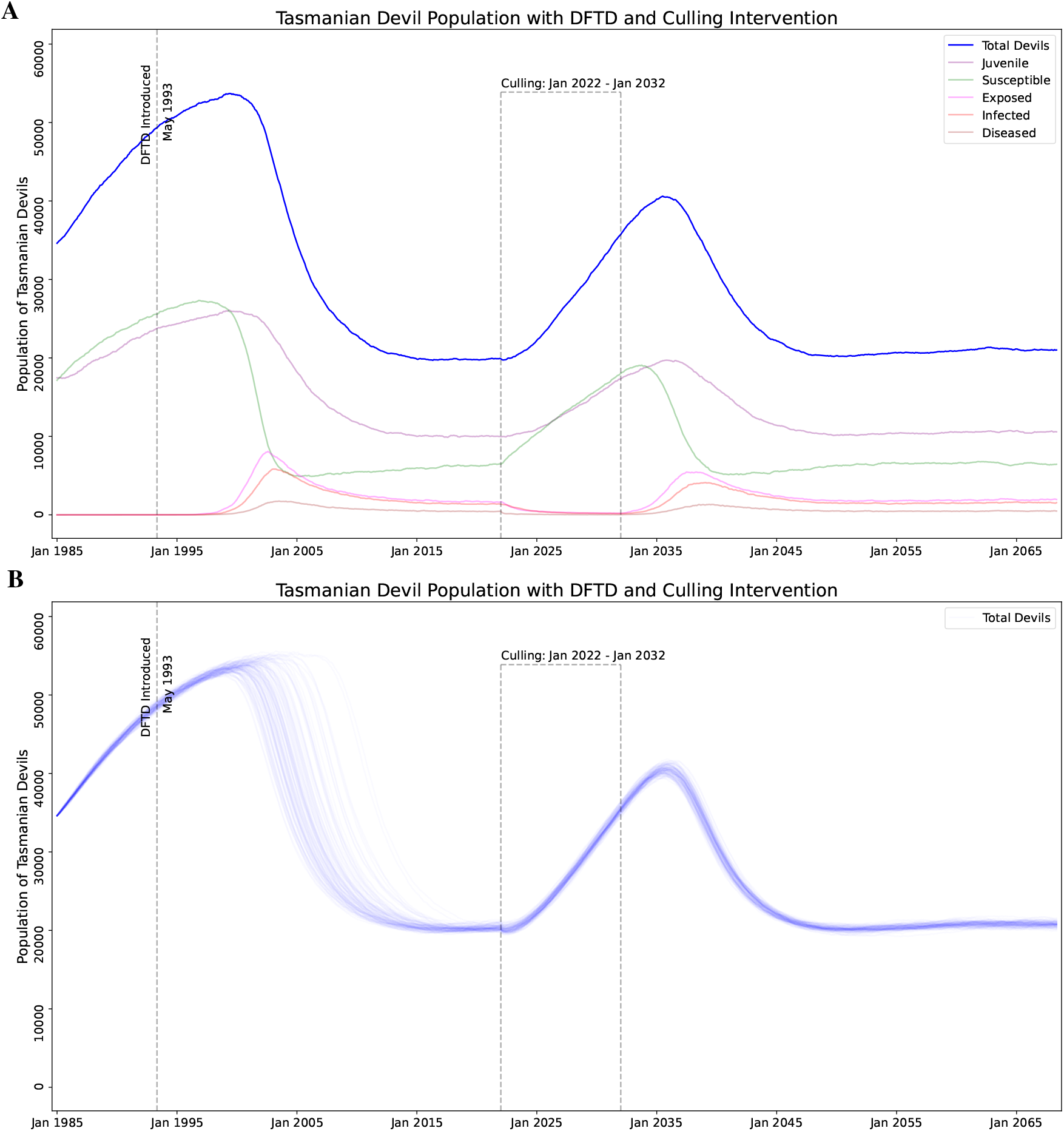
Simulation of DFTD model with a 10 year culling program where 50% of heavily diseased devils removed from the population (Diseased state). **A:** A single stochastic trajectory, showing all the sub-classes. **B:** 100 stochastic trajectories, showing only total devil population.

**Figure 8:**
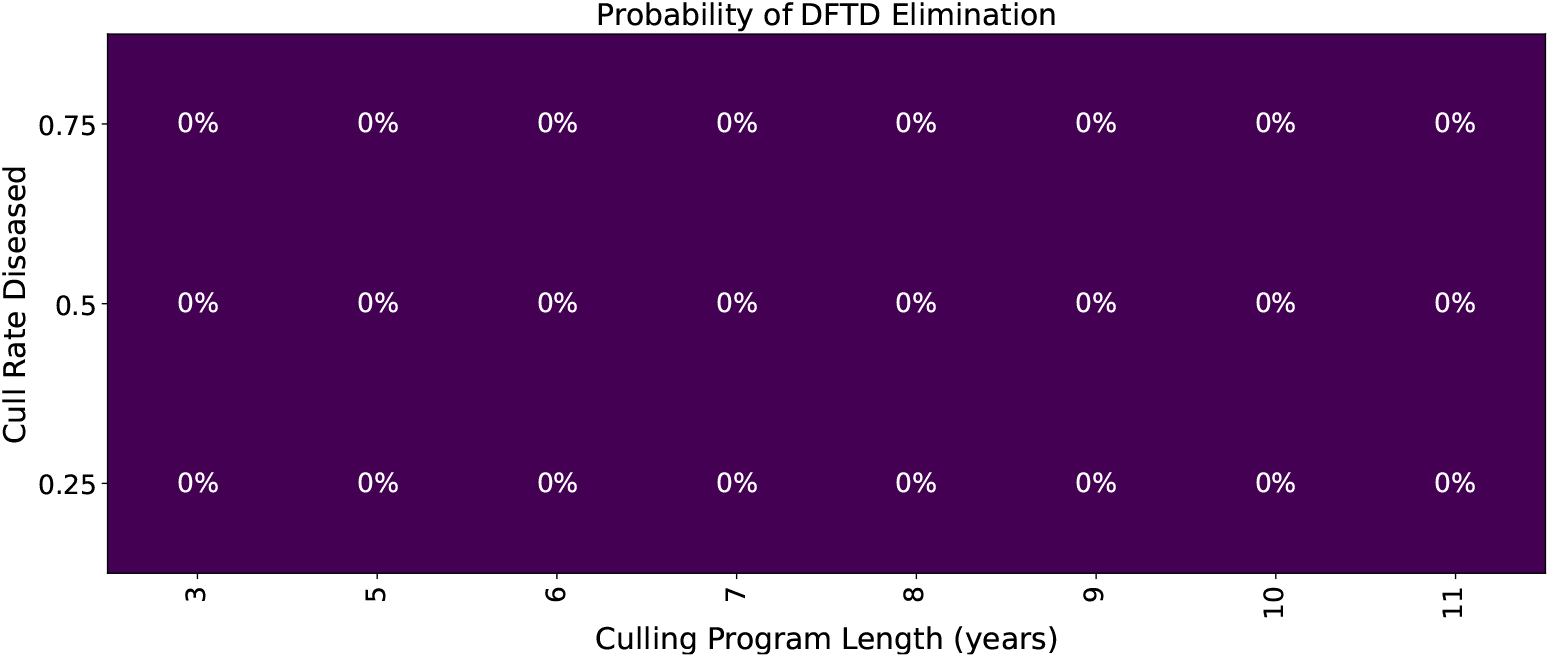
Probability of eliminating DFTD with culling.

### 4.4 Vaccination

Wildlife vaccination against DFTD is still in development but we consider what efficacy of vaccine is needed to be productive as well as how long and how frequent a vaccine campaign would be needed to allow devil populations to recover (Owen and Siddle, 2019; Tovar et al., 2017; Flies et al., 2020). Figure 9 shows the model diagram with the addition of the Vaccinated state, equations (12-14) and Figure 9 shows the additional interactions, and table 4 shows the description and ranges of the vaccination parameters. The model considers the vaccination to be administered from 1-12 times per year for 1-10 years with *P*_*vaccination*_ proportion of the devils receiving the vaccine. The vaccine has an efficacy rate of 1 − *V*_*fail*_, therefore Vaccinated devils become Exposed devils after interacting with an Infected or Diseased devil at reduced infection rates *I*_*infected*_*V*_*fail*_ and *I*_*diseased*_*V*_*fail*_ respectively. Figure 10 shows the model with a ten year vaccination program implemented. A successful campaign where devil populations increase to pre-DFTD populations post the end of the campaign is shown in Figure 10A. This successful campaign assumes vaccine with only 40% efficacy (60% failure rate) that is delivered to 80% of the devil population, six times per year. A campaign that temporarily increases devil populations but does not eliminate the disease and the population decreases back to the endemic steady state is shown in Figure 10B. This campaign assumes the same parameters as the successful campaign but reduces vaccine delivery to only two times per year.

**Table 4:** Parameters for vaccination intervention

| Parameter | Range | Description |
| --- | --- | --- |
| $P_{Vaccination}$ | $0.0 \rightarrow 1.0$ | Fraction of Adult devils vaccinated on each drop |
| $V_{Fail}$ | $0.0 \rightarrow 1.0$ | Probability of vaccine failure |
| Program Length | 1+ | Duration in years of the vaccination program |
| Program Frequency | $1 \rightarrow 12$ | Number of vaccination drops per year |

**Figure 9:**
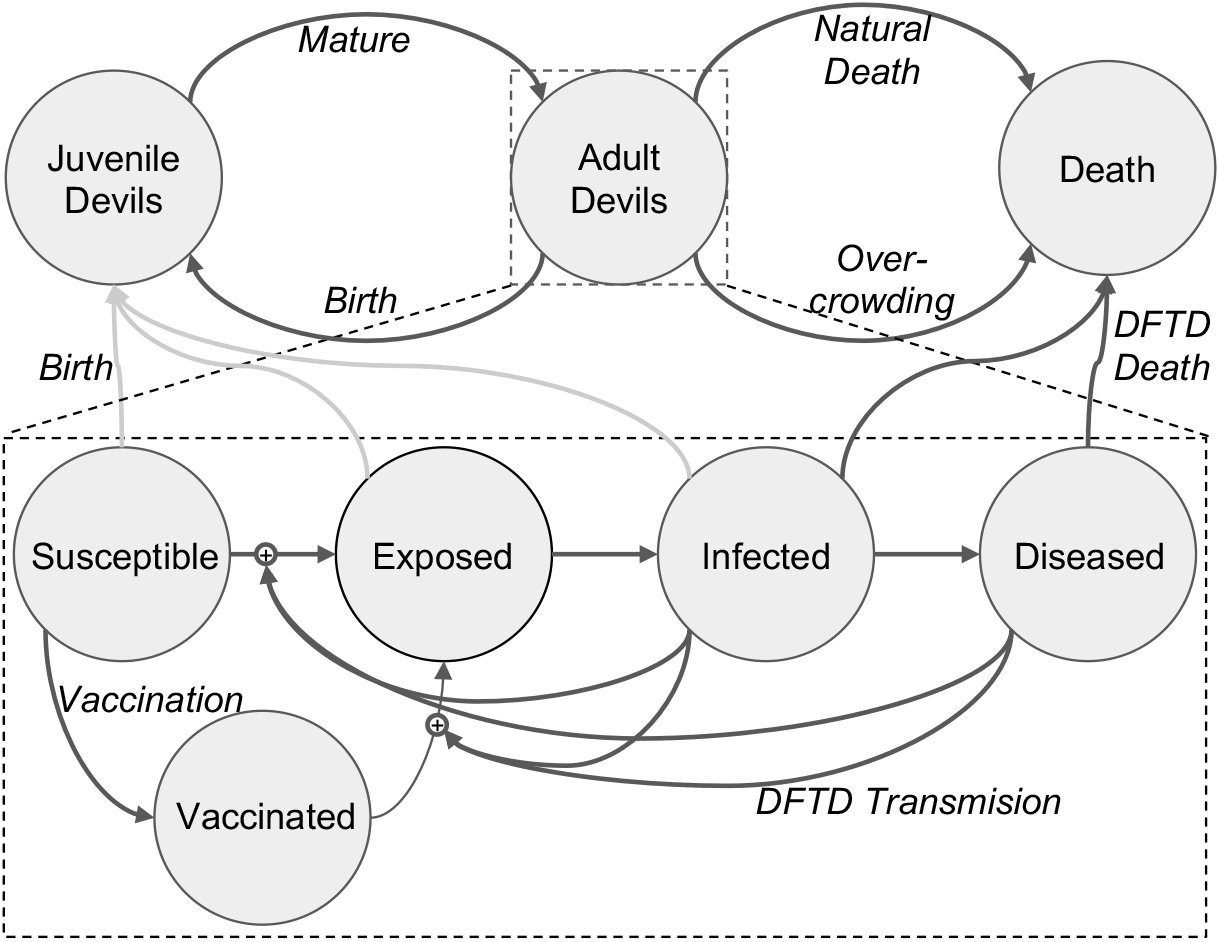
Diagram of our model with the Vaccinated state. We sub-divide “Adult Devil” into five stages of the disease, adding Vaccinated as a subset of the Susceptible population. This state has a lower rate of becoming Exposed given contact with an Infected or Diseased devil.

**Figure 10:**
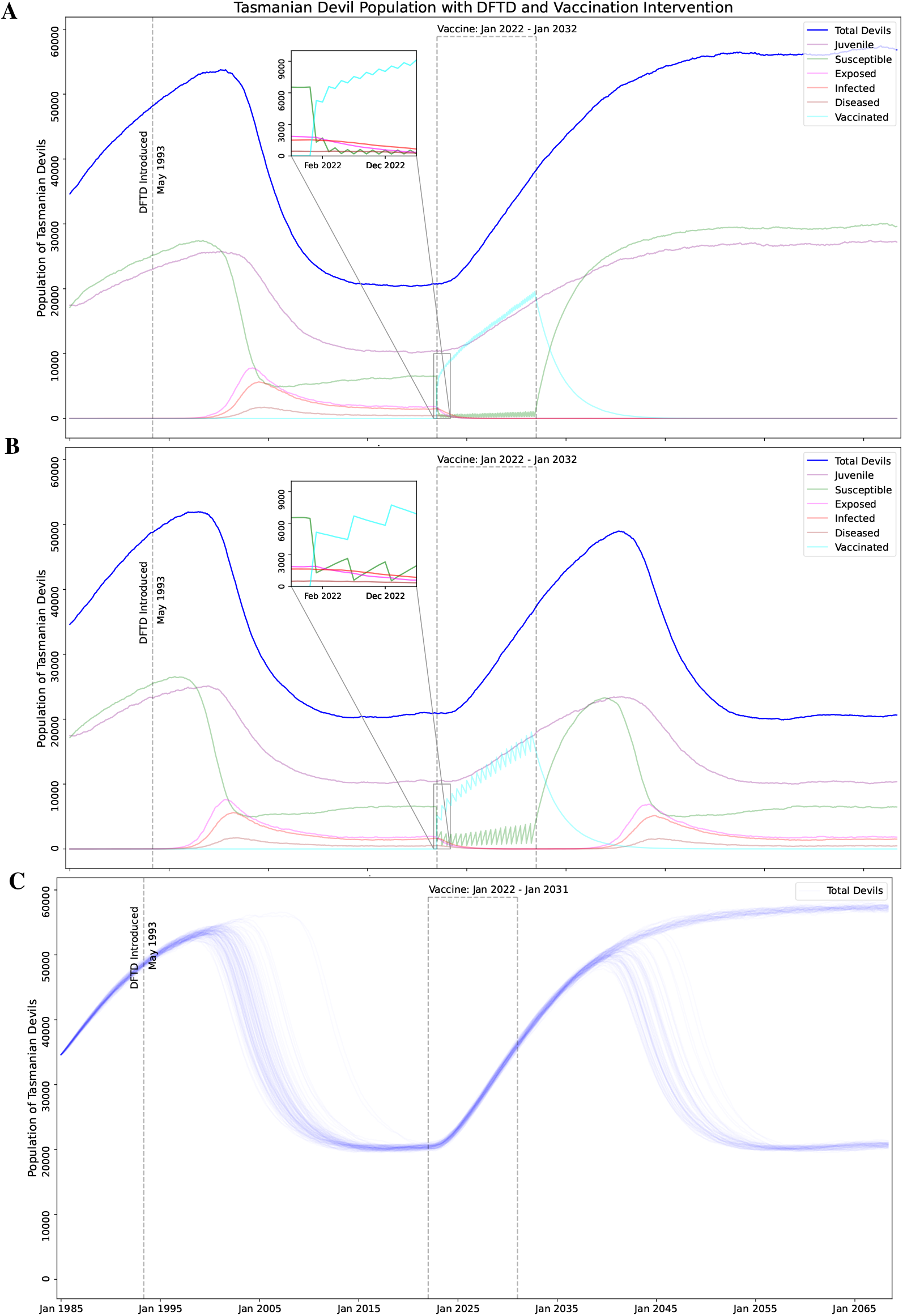
Simulation of DFTD model with a 10 year vaccination program. **A:** A program that successfully eliminates DFTD, using the parameters: 60% vaccine failure rate, delivered to 80% of the population 6 times a year. The inset shows the vaccination events, and resulting spikes in the population of Vaccinated and Susceptibilities. **B:** A program that temporarily increases devil populations but does not eliminate DFTD, using the parameters: 60% vaccine failure rate, delivered to 80% of the population 2 times a year. The inset shows the vaccination events with a different frequency. **C:** A plot of 100 stochastic trajectories, showing approximately 50% DFTD elimination. Simulations used the parameters: 40% vaccine failure rate, delivered to 80% of the population 4 times a year for 9 years. This shows how we calculate the percentage effectiveness of a conservation strategy of a specific parameter set, by counting the number of stochastic trajectories where DFTD is eliminated (blue lines ending near the carrying capacity) divided by the total

We note that due to the high transmissibility of DFTD, if the disease is not successfully eliminated, a single disease animal will eventually push the population back into the current endemic state if no other interventions or acquired immunity develops. Therefore we consider the probability of DFTD elimination based on the vaccine program length in years versus the vaccine frequency per year, the proportion of devils who are vaccinated and the vaccine failure rate (1-vaccine efficacy). Figure 11 shows that in all cases, a program length of at least eight years is needed for at least a 50% probability of DFTD elimination. In Figure 11A we assume a low vaccine failure rate of only 10% with 80% of the devils receiving the vaccine. We notice that a vaccine frequency of at least four to six times per year is needed to achieve over a 50% probability of disease elimination. In Figure 11B we also assume a low vaccine failure rate of only 10% with a frequency of four times per year. We note that developing baits that allow the vaccine to be delivered to a higher proportion of the devils is imperative and the more successful the bait drops are, the fewer years the vaccine campaign can be for potential successful DFTD elimination. Finally in Figure 11C we see something interesting about the vaccine failure rate. We note that the probability of disease does not vary greatly in each column, indicating that vaccine failure rate does not play a strong role in disease elimination. A 10% failure rate (bottom row) does not lead to much greater elimination probabilities than a 60% failure rate (top row) with the same length of the vaccine campaign. Therefore our model supports that with with the development of almost any vaccine, efforts put into successful baits in a long term, frequent vaccine campaign can have a high probability of eliminating DFTD.

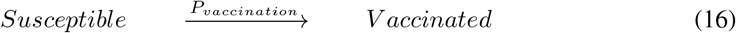

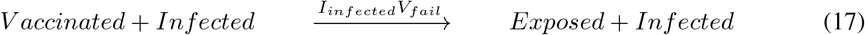

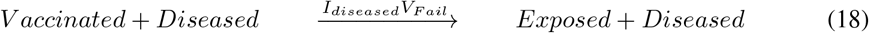

**Figure 11:**
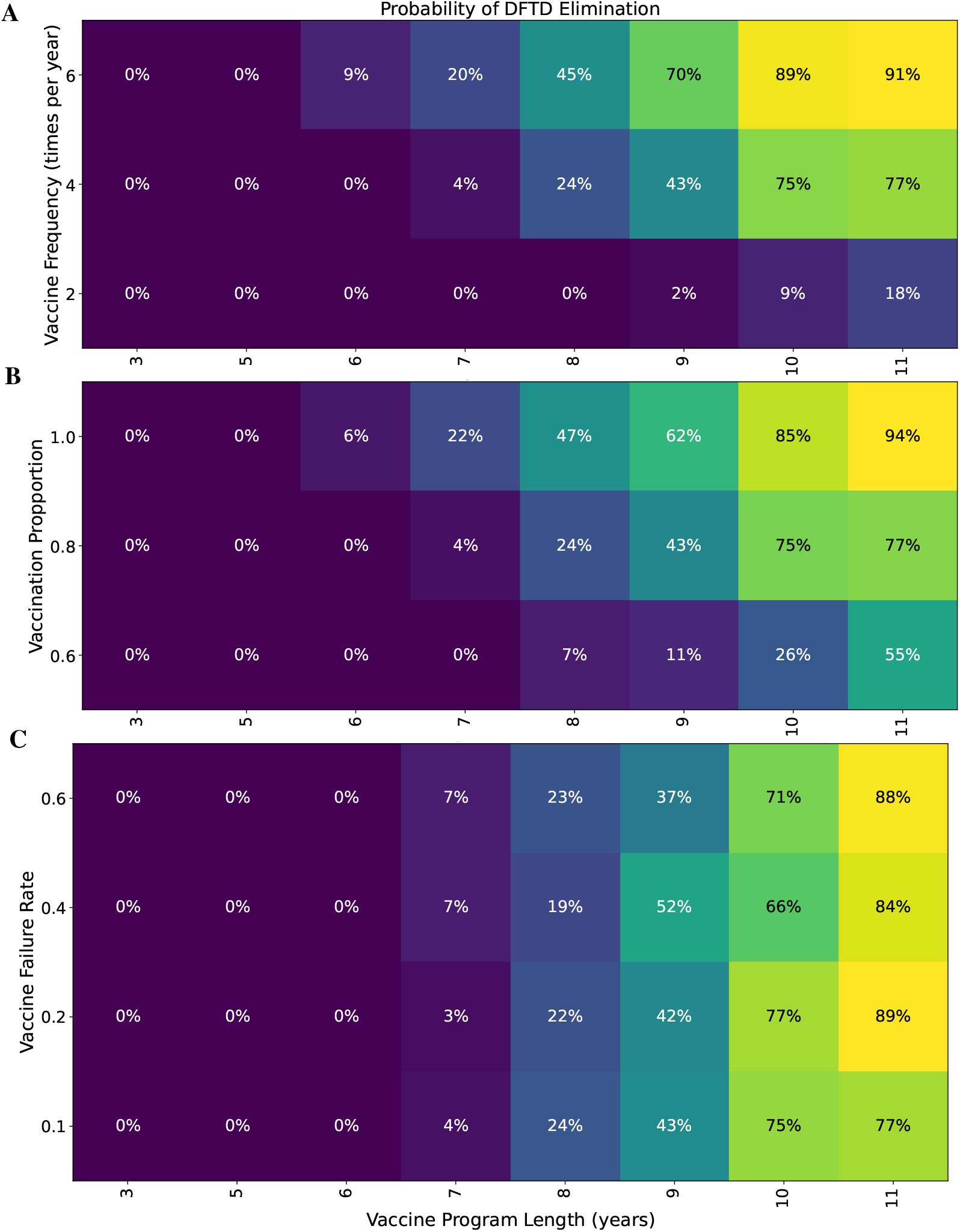
Probability of eliminating DFTD with program length between 3 and 11 years versus **A:** Vaccine frequency (number of times per year) while fixing vaccine failure rate at 0.1, and vaccine proportion at 0.8. **B:** Vaccine proportion while fixing vaccine failure rate at 0.1 and vaccinating frequency 4 times a year. **C:** Vaccine failure rate while fixing vaccinating proportion 0.8 and vaccinating frequency 4 times a year.

Note, that we implement the vaccination process in equation 16 as single event. This mimics the implementation of the proposed bait drop vaccination strategies. Thus our vaccination program has four parameters (as shown in Table 4): *P*_*V accination*_ is the fraction of all wild Tasmanian devil that will be vaccinated by a single drop, *Program Length* is the number of years the conservation will be implements, and *ProgramFrequency* is the number of times per year that bait drops will be performed. We assume a bait drop is a environment wide distribution of vaccine within a short time period (modeled as instantaneous).

### 4.5 Vaccination and Culling Combined

A vaccination campaign can be costly and time consuming, therefore we consider if selective culling of highly diseased animals can reduce the length of a potential vaccine campaign. Animals considered for culling are in the Diseased sub-class meaning they are in the late stages of DFTD, have noticeable visible tumors, and are in the last 3-4 months of their life. While these devils are assumed to not be successfully mating, it is still possible for them to spread the disease to Susceptible devils. Figure 12 shows the effect on the devil population of a program that includes both vaccination and culling. Figure 13 shows that vaccination and culling are more successful than either conservation strategy alone.

**Figure 12:**
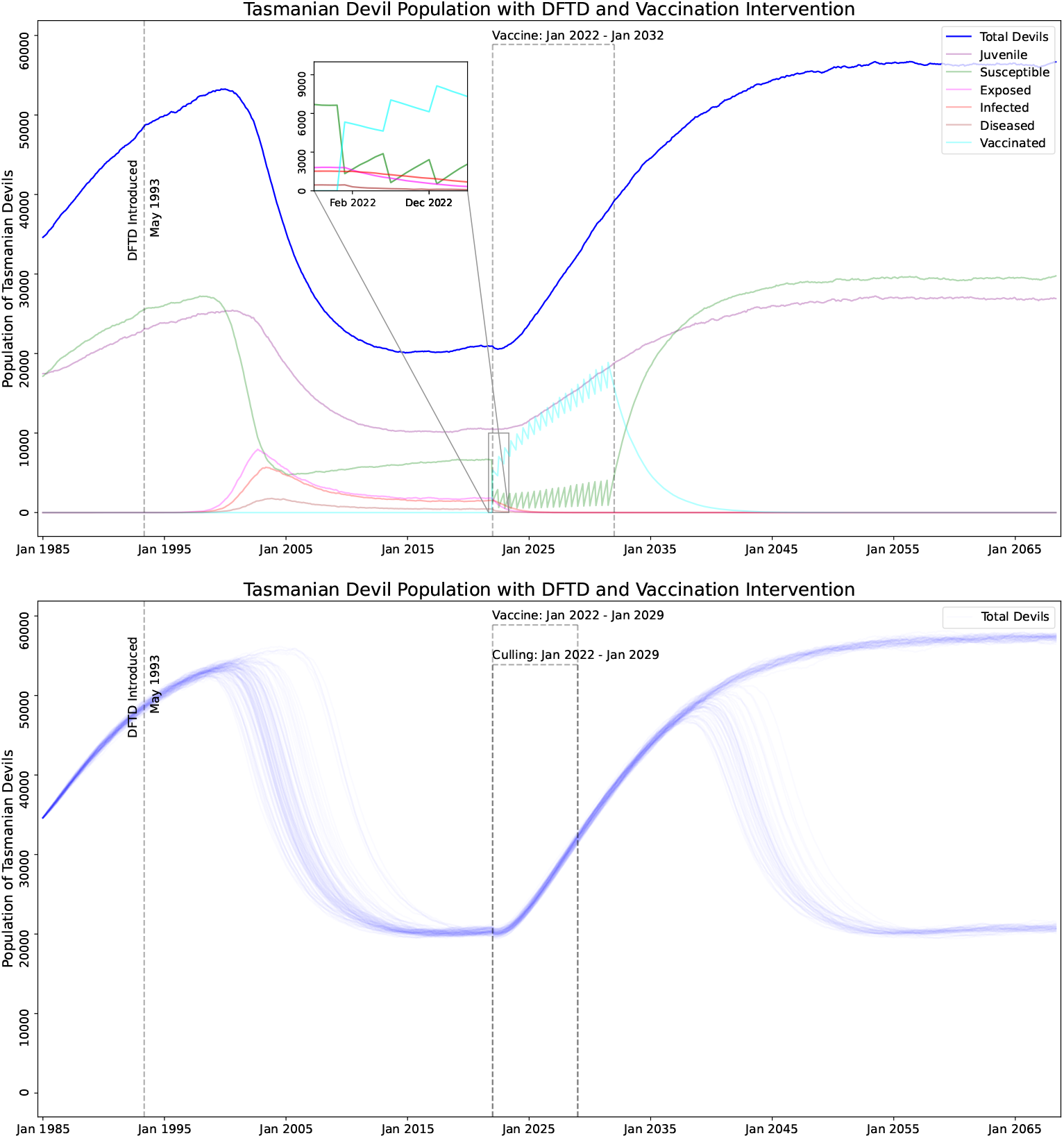
Simulation of DFTD model with both vaccination and culling. **A:** Culling and Vaccination program lengths of 10 years, vaccine failure rate of 0.6, vaccination proportion of 0.8, vaccine frequency of 2 times a year, and a cull rate of 0.5. **B:** Culling and Vaccination program lengths of 7 years, vaccine failure rate of 0.4, vaccination proportion of 0.8, vaccine frequency of 4 times a year, and a cull rate of 0.5.

**Figure 13:**
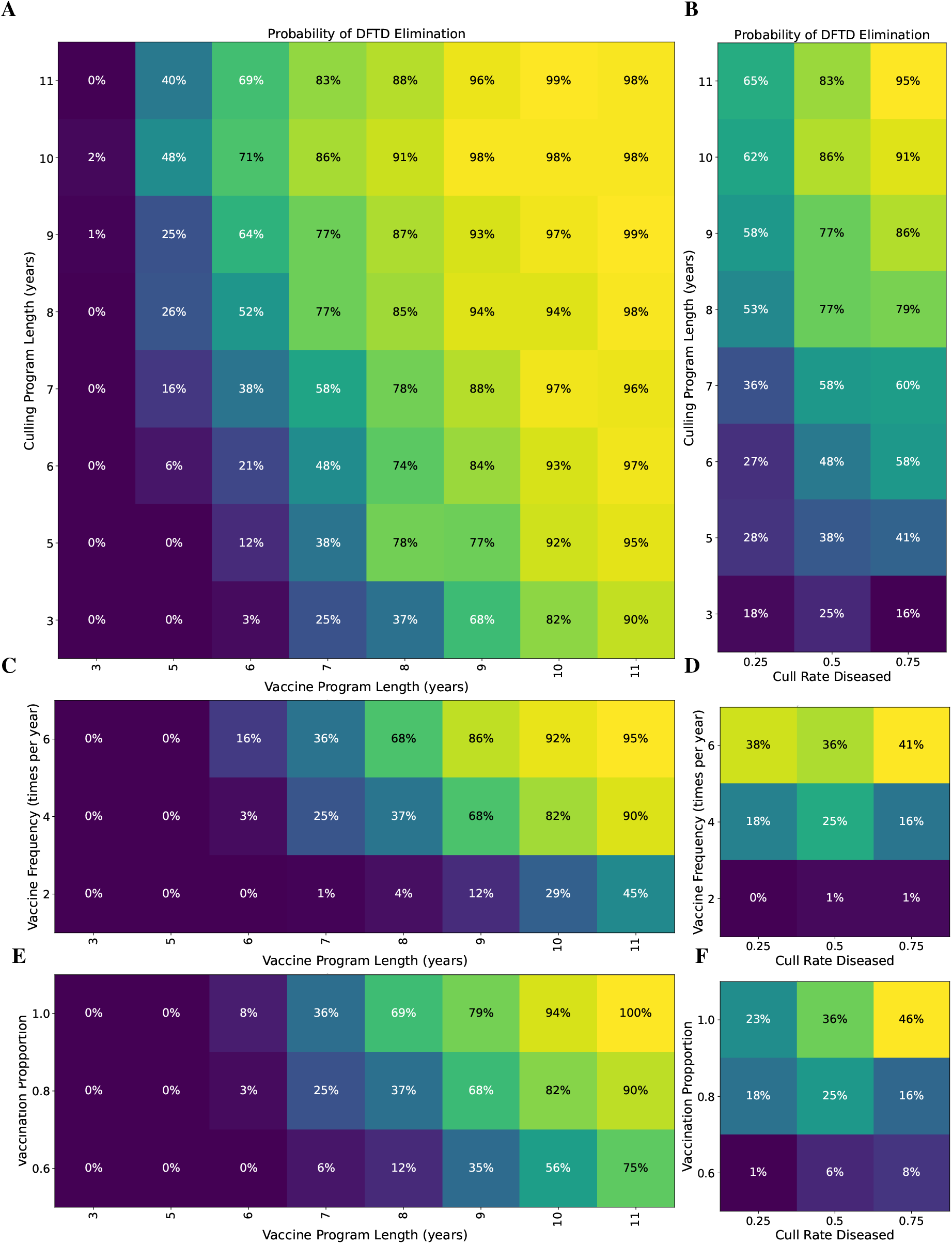
Probability of eliminating DFTD with both vaccination and culling. Parameters held constant (unless specified by the axes) are: vaccine failure rate of 0.4, vaccination proportion of 0.8, vaccination and culling program length of 7 years, vaccine frequency is 4 times a year, and Diseased devil culling rate of 0.5. **A:** Culling Program length versus Vaccination Program length. **B:** Culling Program length versus Diseased devil culling rate. **C:** Vaccine Frequency versus Vaccine Program Length. **D:** Vaccine Frequency versus Diseased devil culling rate. **C:** Vaccine Proportion versus Vaccine Program Length. **D:** Vaccine Proportion versus Diseased devil culling rate.

## 5 Conclusion

In developing a model parameterized based on a 20 years of devil data (Cunningham et al., 2021), we are able to show the potential for DFTD interventions to successfully eliminate the disease. While there is limited evidence that devils are developing natural immunity to the disease, elimination of the disease and population recovery can be supported by a multi-year vaccine campaign, potentially combined with selective culling of highly diseased animals. Our model supports that the devils are currently in the endemic phase and may stay at their current population levels, but with potential changes to the age structure of the population, current devil populations may not be sustainable. A devil population of primarily juveniles, as our model shows the population will become, is problematic and unlikely to lead to a long term healthy population. Therefore, lacking strong evidence for the development of a high level of natural immunity, our model supports vaccine intervention when a viable vaccine become available.

Our simulation results can be interpreted to make a few conclusions. Parameter sweeps of different vaccination strategies show that the results are less sensitive to vaccine failure rate, i.e. decreasing vaccine failure rate does not necessarily increase elimination probability (Figure 11C). Our results show that focusing on vaccine distribution by increasing vaccination frequency and vaccination proportion show increasing chance of elimination (Figure 11A and B). However, for a vaccination program alone results will likely take at least 9 years (results >=50% in figure 11). The timeline for DFTD eradication can be accelerated by combining a vaccination program with a culling program, with possible elimination in 7 years (results >= 50% in figure 13A). It may be beneficial to the long term health of the Tasmanian Devil population to explore some of these options.

## 6 Methods

Our simulations were performed using the software package GillesPy2 (Drawert et al., 2019) (a complete rewrite from the original (Abel et al., 2016)). GillesPy2 is part of the StochSS(Drawert et al., 2016) suite of software (Drawert et al., 2018; Jiang et al., 2021). Our simulation methodology was of discrete stochastic simulation, of the style of the Stochastic Simulation Algorithm (SSA) (Gillespie, 1976, 1977) (sometimes known as the Gillespie method). Specifically, we use the Tau-Hybrid solver with events, which utilizes the Tau-Leaping (Gillespie, 2001) method to simulate the stochastic reaction system.

### 6.1 How to reproduce this work

The modeling, simulation, and analysys were all done with StochSS (Jiang et al., 2021). All of our models and analysis code is hosted on github, and will be posted publicly upon acceptance.

### 6.2 Model Parameterization

We found our set of parameters (see Table 1) using a combination of expert knowledge and a computational technique, similar to the method outlined in (Singh et al., 2020). See Figure 14 for plots showing the sensitivity of the parameter space near the parameter points shown in Table 1.

**Figure 14:**
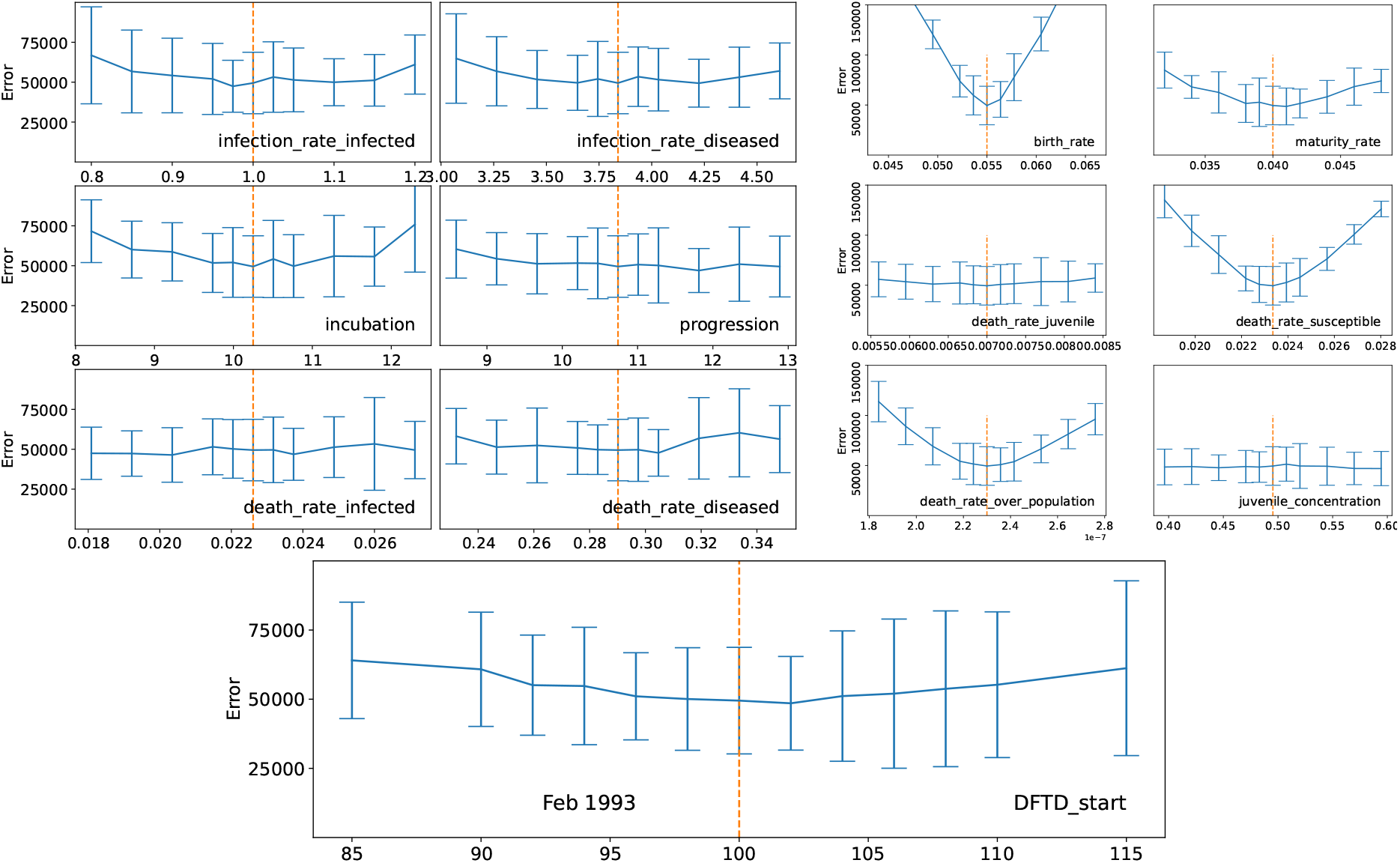
Parameter Space Exploration, plots show the error (L2 norm distance metric) between simulated trajectories and data. Vertical dashed line show selected model parameters (see table 1). Each parameter point was simulated with 1000 stochastic trajectories, the distance to the observed data was calculated for each of trajectories, the mean and standard deviation of that data is shown in the figure.

## 7 Acknowledgements

We acknowledge funding from NIBIB Award 2-R01-EB014877-04A1. No official position or official endorsement should be inferred.

We would like to thank Andy Flies of the University of Tasmania for his feedback and input into this manuscript.

